# Deep representation learning for domain adaptatable classification of infrared spectral imaging data

**DOI:** 10.1101/584227

**Authors:** Arne P. Raulf, Joshua Butke, Claus Küpper, Frederik Großerueschkamp, Klaus Gerwert, Axel Mosig

## Abstract

**Motivation:** Applying infrared microscopy in the context of tissue diagnostics heavily relies on computationally preprocessing the infrared pixel spectra that constitute an infrared microscopic image. Existing approaches involve physical models, which are non-linear in nature and lead to classifiers that do not generalize well, e.g. across different types of tissue preparation. Furthermore, existing preprocessing approaches involve iterative procedures that are computationally demanding, so that computation time required for preprocessing does not keep pace with recent progress in infrared microscopes which can capture whole-slide images within minutes.

**Results:** We investigate the application of stacked contractive autoencoders as an unsupervised approach to preprocess infrared microscopic pixel spectra, followed by supervised fine-tuning to obtain neural networks that can reliably resolve tissue structure. To validate the robustness of the resulting classifier, we demonstrate that a network trained on embedded tissue can be transferred to classify fresh frozen tissue. The features obtained from unsupervised pretraining thus generalize across the large spectral differences between embedded and fresh frozen tissue, where under previous approaches seperate classifiers had to be trained from scratch.

**Availability:** Our implementation can be downloaded from https://github.com/arnrau/SCAE_IR_Spectral_Imaging

## 1 INTRODUCTION

In recent years, the application of label-free infrared microscopy to histopathological tissue samples has paved the way for *spectral histopathology* [3, 11], which has proven to be a reliable approach to assess the disease status of histological sections. Infrared microscopy measures samples with a resolution of few *μ*m and provides an infrared spectrum representing the biochemical tissue status at each pixel location. It has been shown that the pixel spectra obtained from infrared microscopes are highly representative for different tissue components as well as for disease status. As illustrated in Fig. 1, this allows supervised classifiers to infer the tissue component or disease status from an infrared pixel spectrum, which has proven successful for several types of cancer ranging from colon carcinoma [11, 13] to lung [3, 7] and bladder [8] cancer.

It is commonly observed that besides biomedically relevant molecular signatures, data obtained from highly sensitive bioanalytical techniques contain technological or biological artifacts, background signal and other confounders that mask those features that are relevant towards disease status. For infrared microscopy, such background artifacts are particularly severe and complex in nature [1], and several approaches have been proprosed to disentangle them from diagnostically relevant signals. The bandwidth of proposed approaches range from methods based on physical models [1, 12, 18] to statistical approaches utilizing principal components [15]. While these methods have contributed to the successful application of spectral histoapthology in clinical studies, preprocessing infrared spectra remains subject of active investigation in the community [12].

Our contribution breaks with those existing approaches and takes a machine learning perspective on the problem. To elaborate, preprocessing infrared spectra can be viewed as a *representation learning* [2] problem: raw infrared spectra are difficult to classify, so that they need to be transformed into a representation that is more accessible for classification or interpretation. Recent progress on learning such representations in an unsupervised manner suggests that the resulting representations are often more suitable for classification than previous, problem domain specific “feature engineered” representations [2]. The success of unsupervised representation learning is often coupled with the availability of large amounts of data, which are commonly accessible in spectral histopathology. Even a single image commonly contains tens of millions of spectra [11] that can be measured within minutes [14]. In other words, the substantial recent progress in the field of representation learning bears great promises for spectral histopathology that we investigate in this contribution.

**Fig. 1.**
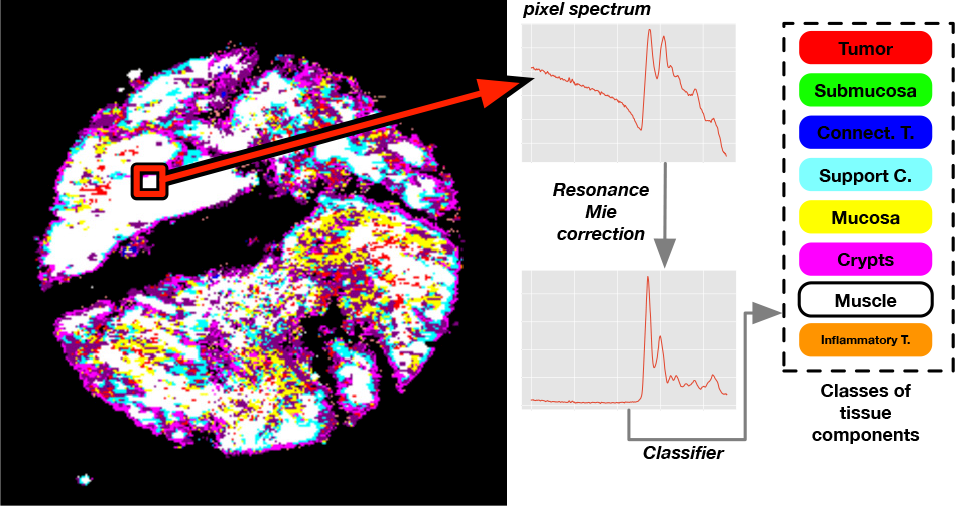
Principle of spectral histopathology: The infrared spectrum from each pixel position is preprocessed using a physical model and subsequently classified into the respective tissue component. Our newly proposed approach aims to use deep neural networks to classify the uncorrected spectrum, where preprocessing is subsituted with an unsupervised deep representation learning step.

The aims of our present contribution go beyond the mere assessment of recent progress in representation learning to spectral histopathology. Specifically, we propose an approach to assess the robustness of a learned representation. We achieve this by transfering a classifier that is based on a learned representation to a related domain. In our specific case, we transfer a classifier trained on data obtained from formalin-fixed tissue to data from fresh frozen tissue, which is accompanied by significant changes in the infrared spectra. If the classifier turns out to be transferable, the learned representation can be considered highly robust against sample variability and heterogeneity. As robustness is of predominant importance when classifying biomedical sample material, we consider the domain transfer approach as a major contribution of our present paper, which can be employed to assess the robustness of not just infrared spectral classifiers, but also of classifiers for data obtained from other bioanalytical techniques.

## 2 BACKGROUND

### Spectral Histopathology

In order to assign disease relevant classes to infrared microscopic pixel spectra, different studies in spectral histopathology have employed a range of different classification approaches. One common approach [11, 7, 3] is to use pixel spectra classifiers to obtain a segmentation of the tissue sample into different physiologically or pathologically relevant components. The segmented image then serves as a basis for a diagnostic characterization, for instance by determining the relative abundance of cancerous or otherwise disease relevant pixels [23]. Some studies [11, 3] suggest that resolving other tissue components along with the distinction into pathological vs. healthy regions is helpful or even neccessary to reliably characterize the disease status. Remarkably, all aforementioned infrared microscopy based studies involve preprocessing of the spectra, typically in the form of physical models that either remove resonance Mie scattering [1] or dispersion artifacts [18].

Until recently, most spectral histopathology studies utilized *Fourier transform infrared* (FTIR) microscopes, where the infrared spectrum is derived by Fourier transforming the signal obtained from an interferometer. Very recently, FTIR microscopy has been challenged by quantum-cascade laser (QCL) microscopes, where the spectrum is obtained from frequency tunable quantum cascade lasers. QCL-based microscopes exceed the measurement speed of FTIR-based systems by almost two orders of magnitude [8], so that infrared images of complete slides of histological sections can be captured within minutes. At the same time, however, the infrared spectrum is limited to a smaller spectral range, which in particular affects the spectral baseline that is important for resonance Mie correction.

### Representation learning and Domain Adaptation

Our work is motivated by groundbreaking progress in the field of representation learning surveyed in [2], which gained much of its momentum from related breakthroughs in the field of deep neural networks and convolutional neural networks [17].

### Transfer learning

An essential question arising from the high accuracies often achieved by extremely parameter rich deep neural networks is whether the network is overfitting the training data rather than having identified truly discriminative features between the classes during supervised training. Observing strong validation measures in conventional cross-validation schemes is certainly a neccessary, but not sufficient criterion. A stronger criterion will be to test how the classifiers perform on previously unseen types of data. In the domain of clinical data, one can identify several levels of what constitutes previously unseen. In [9], it was suggested that validation should be performed at the highest possible level of replication in order to avoid overfitting. In clinical studies, different levels of replication are conceivable such as changes in the measurement device, changes in sample preparation, or even multi-center studies [24].

The concept of robustness is closely related to the *transferability* of classification models: If a classifier generalizes when trained on one specific task, it should be feasible to further generalize classification towards a second task similar to the first task. In a related contribution [10], it was shown that model transfer significantly improves classification of Raman microscopic images across four different microscopes. We investigate transferability in a somewhat broader setting: First, we train a deep neural network on formalin fixed parrafin embedded (FFPE, henceforth referred to as *embedded*) histopathological samples of colon tissue. Then, we investigate a transfer of this classifier to infrared images of fresh frozen tissue samples (henceforth referred to as *fresh tissue*). In related previous studies [13], classifiers were built independently for embedded and fresh tissue, respectively, as model transfer appeared infeasible. In our contribution, we assess transfer learning approaches [16] to facilitate the transfer of an FFPE trained neural network to fresh tissue.

## 3 APPROACH

Our computational approach is summarized in Fig. 2. It is based on first obtaining a gold standard segmentation based on conventionally corrected spectra classified by a conventional, previously established classifier. The only purpose of this pre-segmentation is to obtain a sufficient amount of training data for the second *deep learning* stage. The deep learning stage is in turn divided into two steps: An unsupervised *pre-training* is succeeded by supervised fine-tuning into the final network **pt-MLP**.

As it has been demonstrated [4], these neural network based approaches largely benefit from the availability of large amounts of data. This sets apart our approach in a fundamental way from previous approaches, which rely on manual annotations. As there are inherent difficulties in obtaining suitable annotations on larger numbers of histopathological samples, conventional approaches are favourable towards classifiers that generalize well on small amounts of training data. In this sense, our present contribution aims to establish neural network based approaches that scale with the large amounts of data that are typically available in clinical studies involving infrared microscopy.

Beyond the accuracy of classifier **pt-MLP**, a key question to be assessed is whether **pt-MLP** generalizes well to unseen data sets, or whether it rather overfits the training data. In order to assess the capability of **pt-MLP** to generalize, we perform *transfer learning*. Specifically, we employ a second set of colon cancer related tissue samples. This set of tissue samples and the image spectra obtained from it differ substantially from the first set: first, this data set has been acquired from fresh tissue rather than paraffin embedded tissue. Second, the samples were obtained as full sections rather than as tissue microarrays. The substantial differences in the image spectra are illustrated in Supplementary Fig. 1. A gold standard pre-segmentation for producing training data was applied in a similar fashion as for the first data set.

For convenience, Supplementary Table 1 provides an overview of the different supervised classifiers and their role in this study.

## 4 METHODS

### Sample Material

We employed two datasets of infrared images of histopathological samples related to colon cancer that have been investigated in previous studies.

The first dataset from [13] consists of infrared microscopic images of embedded tissue microarray (TMA) samples. We employed two such TMA slides purchased from *US Biomax Inc., MD, USA*, referred to by their IDs *CO1002b* and *CO722*, respectively, each of which consists of 100 circular spots of tissue sample. Each spot has a diameter of roughly 1 mm. Along with this data set, we also utilized the random forest classifier established previously in [13], which classifies resonance Mie corrected infrared pixel spectra into 19 different classes representing 13 types of tissue components and subclasses, as depicted in Supplementary Fig. 3. This random forest classifier is referred to as classifier **RF** in Fig. 2 and throughout the rest of this manuscript. We will refer to this dataset as the *FFPE dataset*. The FFPE dataset was available and utilized both in the form of uncorrected spectra and resonance Mie corrected spectra preprocessed by the approach from [1]. The FFPE data set was subdivided into three parts, one part for unsupervised pre-training, the second subset for supervised fine-tuning, and the third subset was withheld for validation. The validation subset was strictly seperated from the other two parts and was neither involved in pre-training nor in supervised learning.

Our second dataset was acquired from fresh-frozen histopathological colon tissue samples. Each sample is represented by one infrared image covering a whole tissue section roughly 2 cm^2^ in size. The dataset involves three such whole-slide images. For this data set, we utilized a corresponding classifier **RF2** that was established previously. The training data for classifier **RF2** have been obtained fully independent of classifier **RF**. Throughout this manuscript, we will refer to this dataset as the *fresh tissue dataset*.

In both data sets, the FFPE as well as the fresh data set, pixel spectra with low signal intensity were filtered out and marked as background based on previously described practice [11, 13]. The filtering is performed on uncorrected raw spectra, so that it does not affect our approach being independent from the physical model based resonance Mie correction.

### Obtaining ground truth segmentations

In order to obtain uncorrected spectra with labels for supervised training, we applied classifier **RF** which was previsouly established in [13] to all resonance Mie corrected spectra from the FFPE dataset. Since the uncorrected counterpart is immediately available for each corrected pixel spectrum, this allowed us to assign a class to each uncorrected spectrum. As illustrated in Fig. 2, we use the resulting assignment between uncorrected spectra and tissue components as ground truth for the training data set for our deep neural networks. Note that the classification outcome of **RF** cannot be assumed to be 100% correct on a per-pixel basis and the assignment thus obtained constitutes a *gold standard* in the sense of the best-possible per-pixel annotation rather than a ground truth. Despite this somewhat curtailing factor, we will refer to the training labels obtained from **RF** as ground truth.

### Learning regularized representations through autoencoders

Formally, an autoencoder is constituted by a neural network that represents a mapping *A*: ℝ^*d*^ → ℝ^*d*^, i.e., a network whose input and output layers consist of *d* neurons each. In its most basic form, an autoencoder involves one hidden layer with *M* < *d* neurons. A sequence of (*d, M*_1_), (*M*_1_, *M*_2_), …, (*M*_*K*−1_, *M*_*K*_) autoencoders can be cascaded in a straighforward manner as illustrated in Supplementary Fig. 2 and detailed in Supplement S.1. In [21], such *stacked autoencoders* have been proposed and successfully established as highly effective regularizers on several data sets.

Our stacked autoencoder preprocesses spectra represented as a vector featuring absorbances at *d* = 450 many wavenumbers. The stacked autoencoder involved six hidden layers of sizes *M*_1_, …, *M*_6_ = 450; 900; 450; 100; 100; 100. The stacked autonecoder was trained in an unsupervised fashion following [20] on 2, 220, 000 spectra obtained from 25 spots of TMA slide *CO1002b*. Each autoencoder was initialized following [6].

### Supervised finetuning for classification of pixel spectra

We followed the approach by Rifai *et al* [17] and employed the autoencoder described in Section 4 as an unsupervised pretraining procedure to improve a subsequent supervised learning step. To this end, a *softmax* output layer with one output neuron for each of the 19 classes was added to the six encoding layers of the stacked autoencoder. The resulting network topology was initialized with the parameters of the stacked autonecoder for the first six layers, and random values for the input weights of the output layer. The last three hidden layers were treated as drop out layers [19] with a drop out rate of 50%. This network was trained on the ground truth provided by classifier **RF** as shown in Fig. 2 to obtain supervised classifier **pt-MLP** using *RMSProp* optimization running for 15, 000 epochs.

As a reference to assess the performance of **pt-MLP**, we trained a conventional multilayer perceptron **MLP** based on the same topology as network **pt-MLP**. As in classifier **pt-MLP**, the last three hidden layers were implemented as drop out layers with a dropout rate of 50%. We initialized all parameters in the network randomly and trained it against the same ground truth using *RMSprop* for optimization running 15, 000 iterations.

### Transfer learning for domain adaptation

In order to assess the generalization capability of the unsupervised pretraining procedure and the resulting classifier **pt-MLP**, we performed transfer learning from FFPE to fresh tissue samples. Ground truth on fresh tissue for transfer learning was obtained from a previously established classifier **RF2**, which was trained on resonance Mie corrected spectra in fresh tissue. To adapt to the different classes of tissue components annotated in FFPE vs. fresh tissue (see Supplementary Fig. 3), the output layer was substituted by a randomly initialized softmax layer. The transfer learning approach is illustrated in Supplementary Fig. 5.

We divided data set *Fresh 1* into a trainig data set and a test data set for transfer learning, and performed 15, 000 epochs of *RMSProp* training. Data sets *Fresh 2* and *Fresh 3* were used for validation. For technical reasons, data set *Fresh 3* was subdivided into data sets *Fresh 3A* and *Fresh 3B* during validation.

Details of determining accuracies for validation are described in Supplement S.2.

## 5 RESULTS

### Pretraining with SCAE and supervised Finetuning

We performed pretraining as described in Section 4 on 2.2 million spectra from the FFPE data set *CO722*. The deep learning classifier **pt-MLP** was obtained by finetuning as described in Section 4 on 1.3 million spectra from data set *CO1002b*.

Fig. 3 demonstrates the generalization capability on a held-back TMA dataset. The per-pixel accuracy of classifier **pt-MLP** reconstructing the ground truth segmentation of classifier **RF** achieved an accuracy of 96%.

### Transfer-Learning Segmentations of Fresh Tissue

To assess the transferability of classifiers from FFPE tissue to fresh tissue, we first visualized spectral differences between FFPE tissue and fresh tissue, which is illustrated for two classes of tissue components in Supplemtary Fig. 1. In particular at the level of uncorrected raw spectra, these differences are substantial, so that applying classifiers trained on FFPE spectra to fresh tissue can be expected to lead to very limited success. To examine this in practice, we applied the FFPE-trained classifier **pt-MLP** to fresh tissue. As ground truth, we used the segmentation obtained from a previously established random forest classifier **RF2**. As shown in Fig. 4, **pt-MLP** performs poorly in identifying the tissue structure, achieving an accuracy of only 53%. Two classes, namely submucosa and muscle, were at least partially detected correctly by **pt-MLP**.

Finally, we performed transfer learning from FFPE tissue spectra to fresh tissue spectra by training classifier **tl-MLP** using **pt-MLP** as a starting point. As training data for transfer learning, we used uncorrected fresh tissue spectra labelled with the output classifier **RF2** as ground truth, as illustrated in Fig. 5. A validation of the result is shown in Figures 4 and Supplementary Fig. 6, demonstrating the high accuracies of 82%, 72% and 80% on data sets *Fresh 2*, *Fresh 3A* and *Fresh 3B*, respectively of the transfer learned classifier **tl-MLP**.

**Fig. 2.**
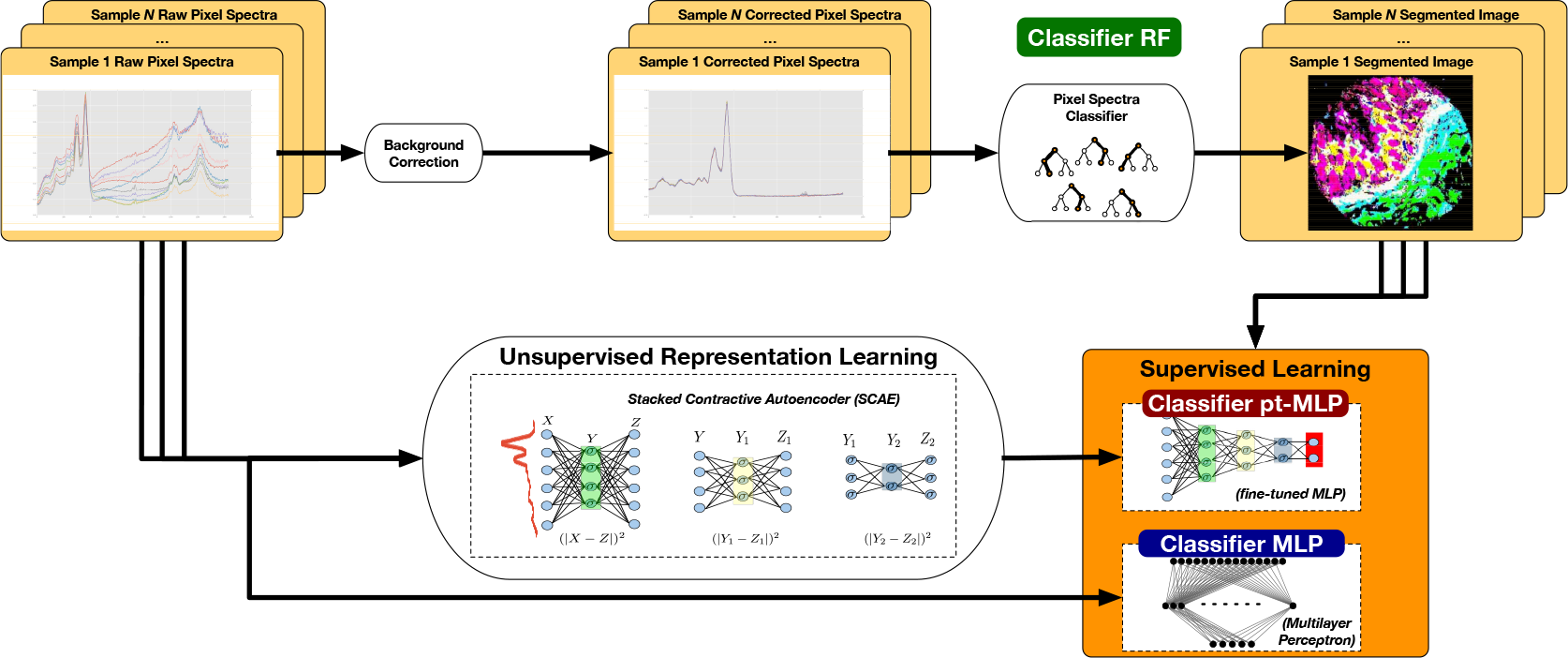
Workflow for obtaining ground truth results, performing unsupervised pre-training, and finally supervised fine-tuning as described in Section 3.

**Fig. 3.**
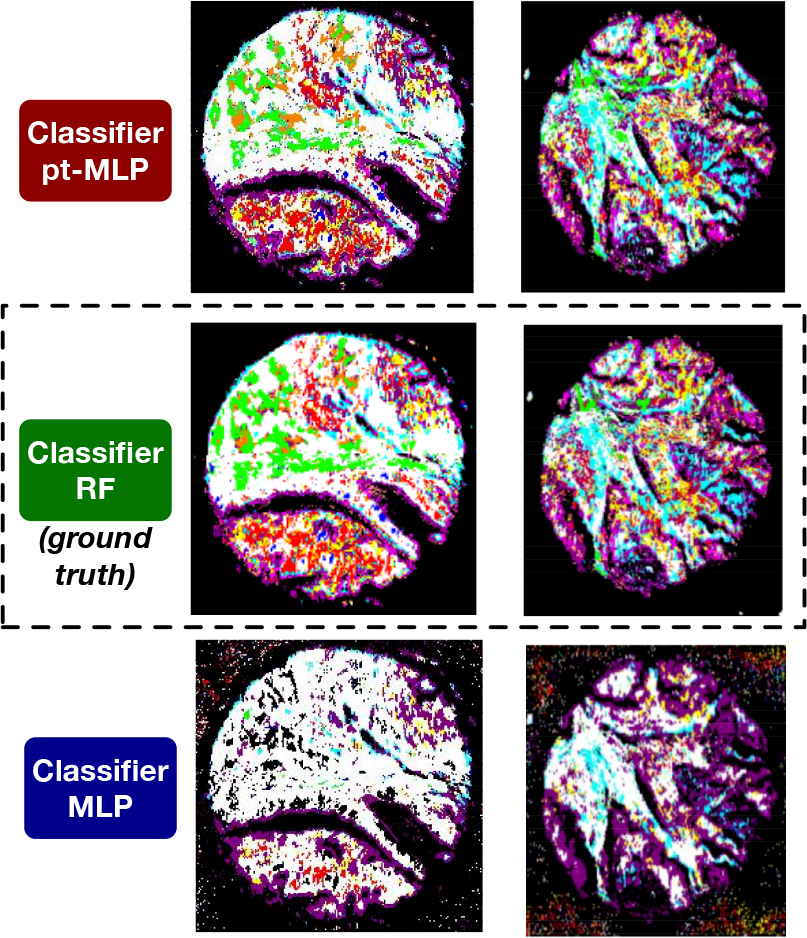
Classification of FTIR raw data of two FFPE-embedded Tissue Microarray spots (TMA) from the fully independent validation data set. The agreement between the pre-trained **pt-MLP** (first row) and the ground truth provided by **RF** (middle row) is very high, and differences recognizable only in small details. The non-pretrained classifier MLP, on the other hand, exhibits systematic misclassification, which is most notably of crypts in the tumor-free spot (right, index color pink) and the tumor and the submucosa class (red and green), which are systematically misclassified as muscle (white) in the tumor spot (left). Classification results of further spots are displayed in Supplementary Fig. 4.

## 6 DISCUSSION

In our study, we demonstrated that unsupervised pre-training of infrared spectra facilitates highly accurate classification of spectral histopathology imaging data. Unsupervised pre-training takes the role of spectral pre-processing, which previously has been tackled on the grounds of physical models in combination with conventional classifiers. Thus, we have demonstrated that a purely data-driven approach can take the role that has previously been taken by a physical model when classifying pixel spectra.

A natural question that arises in this context, and in neural networks in general, is to characterize and interpret what the neural network actually learned. Our results allow the conclusion that the network *implicitely* identified variances that are eliminated by correction procedure underlying the resonance Mie model by Basan *et al.* [1], because classifier **pt-MLP** can reproduce the classification of resonance Mie corrected spectra. This is remarkable since this implies that the network has learned to disentangle the complex interference between molecular spectrum and the scattering artifact, which is neither additive nor linear.

**Fig. 4.**
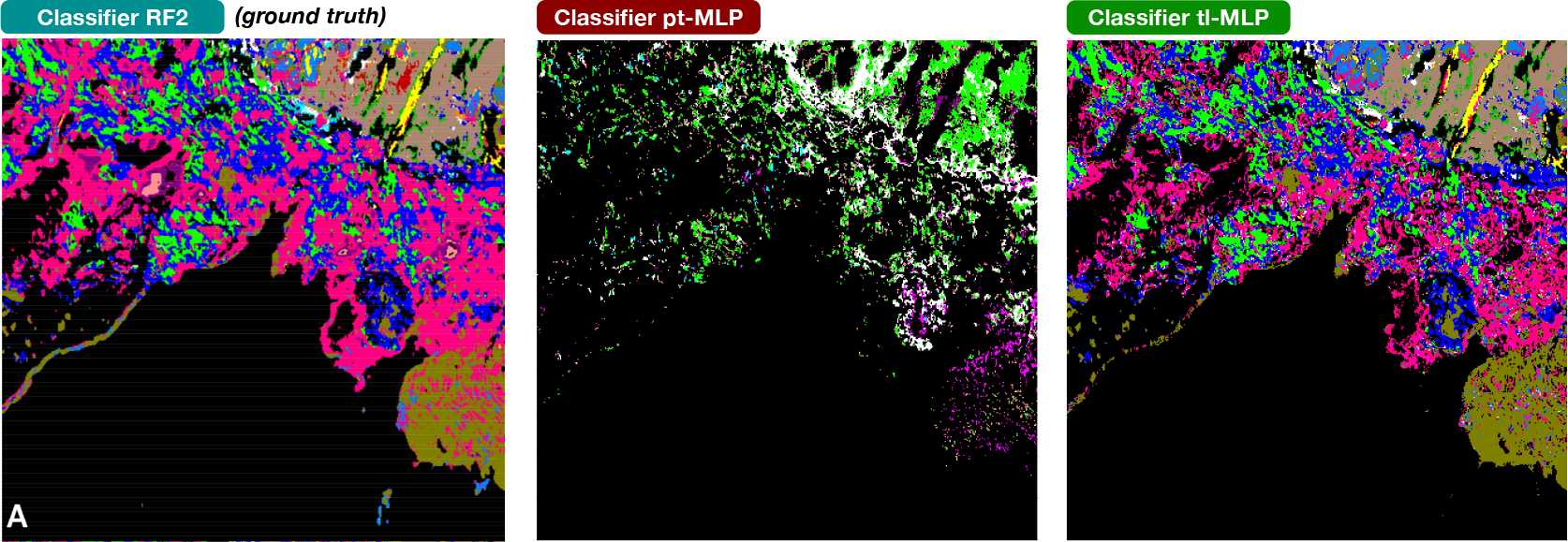
Classification of FTIR raw data with and without transfer learning on independent validation data set *Fresh 2*. *Left:* ground truth obtained from classifier **RF2** on Mie-scattering corrected FTIR-microspectroscopy imaging; *Middle:* prediction obtained from the FFPE-based classifier **pt-MLP**, which fails to identify most of the tissue components in fresh tissue and achieves an accuracy of only 53%. *Right:* prediction of the transfer learned deep learning classifier **tl-MLP**. Results for data set *Fresh 3* are shown in Supplementary Fig. 6.

Since the resonance Mie model is neither explicitely nor implicitely involved in the procedure of training the network, the question arises whether the network may have learned a much more general representation of infrared pixel spectra from histopathological samples. While beyond the scope of our current contribution, this question of model interpretation points to an interesting and relevant new direction for future research, namely to correlate the output of the stacked autoencoder with different physical model-based correction procedures. More specifically, one could investigate how well a spectrum corrected by a physical model can be reconstructed from the representation learned by the neural network. This could be realized by establishing a neural network that learns to approximate a given correction procedure, using the representation learned by the unsupervised pre-training as a starting point. If such a network could reconstruct the result of given physical model-based preprocessing procedure, one would obtain a more *explicit* proof that the network has learned a certain preprocessing function.

As we have argued, there is strong evidence that the stacked autoencoder did learn a meaningful representation of infrared pixel spectra. An obvious next question is how far this representation will *generalize*: how much variance can be added and what type of variance can be added before networks derived from the representation will lose their classification capability. Our approach to determine generalization capability was to investigate the capability of the network to perform *domain adaptation*: If the network can adapt from the domain of embedded tissue to the domain fresh tissue, the underlying representation must be sufficiently abstract and thus generalizable. The high accuracies we observe in the transfer learned networks for fresh tissue strongly indicate that indeed the representation is sufficiently general. It is quite remarkable that despite the moderate effort used for transfer learning – we used the same number of epochs for transfer learning **tl-MLP** as for fine-tuning the original network **pt-MLP** while the last hidden layer of **tl-MLP** had to be fully re-initialized due to the differences in ground truth classes – we obtain very high accuracies.

As the generalization capability is strong enough to adapt an FFPE classifier to fresh tissue, the question emerges how strongly the network will generalize. There is a broad bandwidth of conceivable sources of variance over which it is desirable to obtain a general spectral representation: Besides the variation in tissue type (FFPE vs. fresh) investigated here, there may be variation in the type of microscope (FTIR vs. QCL), variation in the substrate, or variation in the organ of origin (colon vs. tissue form other organs), to mention only a few. A key question to be adressed will be which variation needs to be included in the data for pre-training the stacked autonecoder, and to which variation will the resulting network be capable of generalizing. As we have shown in our work, not all variation needs to be reflected in the pre-training data: Although we did not use any fresh tissue for pre-training, the network turned out to be transferable to fresh tissue anyway. An ultimate goal may be to train an universal preprocessing network that generalizes broadly across the aforementioned sources of variance. We limit our claims of the pre-training performed in the present study for the network to generalize across tissue type, i.e., across FFPE and fresh tissue. This constitutes a major progress over the previously employed physical model based preprocessing, whose generalization capabilities are inherently limited if present at all.

Finally, generalization capability of classifiers against different sources of variance certainly is a key aspect for robust classifiers in life science research in general, and biomarker discovery in particular. As our approach to combine representation learning with domain adaptation involves no assumptions specific to infrared microscopy, this observation bears promise well beyond infrared microscopy. In fact, baseline correction or other forms of preprocessing commonly constitute problems in the analysis of various types of bioanalytical data, ranging from NMR spectroscopy [22], mass spectrometry [5] or Raman spectroscopy [25]. While several different approaches have been proposed for these techniques, it has been commonly observed that preprocessing, in some cases heavily, affects subsequent analysis [5]. On the other hand, the baseline artifacts in other bioanalytical spectra tend to be less complex than artifacts in infrared spectra, and it thus appears reasonable to assume that other types of bioanalytical data can greatly benefit from our purely data-driven unsupervised preprocessing approach using stacked autonecoders.

## 7 CONCLUSION

We have tackled and successfully solved two related problems in spectral histopathology that have not been studied previously, likely because they could not be solved with conventional classifiers: First, we demonstrated that unsupervised pre-training allows to train classifiers that can classify unprocessed raw pixel spectra of infrared microscopic images, and thus may substitute the physical model based preprocessing of infrared image spectra. At the same time, these classifiers are well-established regularizers and thus hold the promise of generalizing stronger and having less tendency towards overfitting. Second, we have demonstrated the transfer of a classifier from the domain of FFPE samples to the domain of fresh tissue, which previously required collecting substantial amounts of new training data and training a classifier from scratch.

## ACKNOWLEDGEMENT

We want to thank Angela Kallenbach-Thieltges for providing the RF classifier for FFPE tissue thin sections. This research was supported by the Protein Research Unit Ruhr within Europe (PURE) funded by the Ministry of Innovation, Science and Research (MIWF) of North-Rhine Westphalia, Germany (grant number: 233-1.08.03.03-031-68079).

## REFERENCES

[1] Paul Bassan, Achim Kohler, Harald Martens, Joe Lee, Hugh J Byrne, Paul Dumas, Ehsan Gazi, Michael Brown, Noel Clarke, and Peter Gardner. Resonant mie scattering (rmies) correction of infrared spectra from highly scattering biological samples. Analyst, 135(2):268–277, 2010.

[2] Yoshua Bengio, Aaron Courville, and Pascal Vincent. Representation learning: A review and new perspectives. IEEE transactions on pattern analysis and machine intelligence, 35(8):1798–1828, 2013.

[3] Benjamin Bird, Milos Miljkovic, Stan Remiszewski, Ali Akalin, Mark Kon, and Max Diem. Infrared spectral histopathology (shp): a novel diagnostic tool for the accurate classification of lung cancer. Laboratory investigation, 92(9):1358, 2012.

[4] Xue-Wen Chen and Xiaotong Lin. Big data deep learning: challenges and perspectives. IEEE access, 2:514–525, 2014.

[5] Pan Du, Warren A Kibbe, and Simon M Lin. Improved peak detection in mass spectrum by incorporating continuous wavelet transform-based pattern matching. Bioinformatics, 22(17):2059–2065, 2006.

[6] Xavier Glorot and Yoshua Bengio. Understanding the difficulty of training deep feedforward neural networks. In Proceedings of the thirteenth international conference on artificial intelligence and statistics, pages 249–256, 2010.

[7] Frederik Großerueschkamp, Angela Kallenbach-Thieltges, Thomas Behrens, Thomas Brüning, Matthias Altmayer, Georgios Stamatis, Dirk Theegarten, and Klaus Gerwert. Marker-free automated histopathological annotation of lung tumour subtypes by ftir imaging. Analyst, 140(7):2114–2120, 2015.

[8] Frederik Großerueschkamp, Thilo Bracht, Hanna C Diehl, Claus Kuepper, Maike Ahrens, Angela Kallenbach-Thieltges, Axel Mosig, Martin Eisenacher, Katrin Marcus, Thomas Behrens, et al. Spatial and molecular resolution of diffuse malignant mesothelioma heterogeneity by integrating label-free ftir imaging, laser capture microdissection and proteomics. Scientific reports, 7:44829, 2017.

[9] Shuxia Guo, Thomas Bocklitz, Ute Neugebauer, and Jürgen Popp. Common mistakes in cross-validating classification models. Analytical Methods, 9(30): 4410–4417, 2017.

[10] Shuxia Guo, Achim Kohler, Boris Zimmermann, Ralf Heinke, Stephan Stoöckel, Petra Roösch, Juürgen Popp, and Thomas Bocklitz. Extended multiplicative signal correction based model transfer for raman spectroscopy in biological applications. Analytical chemistry, 90(16):9787–9795, 2018.

[11] Angela Kallenbach-Thieltges, Frederik Großerüschkamp, Axel Mosig, Max Diem, Andrea Tannapfel, and Klaus Gerwert. Immunohistochemistry, histopathology and infrared spectral histopathology of colon cancer tissue sections. Journal of biophotonics, 6(1):88–100, 2013.

[12] Tatiana Konevskikh, Rozalia Lukacs, and Achim Kohler. An improved algorithm for fast resonant mie scatter correction of infrared spectra of cells and tissues. Journal of biophotonics, 11(1):e201600307, 2018.

[13] Claus Kuepper, Frederik Großerueschkamp, Angela Kallenbach-Thieltges, Axel Mosig, Andrea Tannapfel, and Klaus Gerwert. Label-free classification of colon cancer grading using infrared spectral histopathology. Faraday discussions, 187: 105–118, 2016.

[14] Claus Kuepper, Angela Kallenbach-Thieltges, Hendrik Juette, Andrea Tannapfel, Frederik Großerueschkamp, and Klaus Gerwert. Quantum cascade laser-based infrared microscopy for label-free and automated cancer classification in tissue sections. Scientific reports, 8(1):7717, 2018.

[15] Ellen J Swain Marcsisin, Christina M Uttero, Antonella I Mazur, Miloš Miljković, Benjamin Bird, and Max Diem. Noise adjusted principal component reconstruction to optimize infrared microspectroscopy of individual live cells. Analyst, 137(13):2958–2964, 2012.

[16] Sinno Jialin Pan, Qiang Yang, et al. A survey on transfer learning. IEEE Transactions on knowledge and data engineering, 22(10):1345–1359, 2010.

[17] Salah Rifai, Pascal Vincent, Xavier Muller, Xavier Glorot, and Yoshua Bengio. Contractive auto-encoders: Explicit invariance during feature extraction. In Proceedings of the 28th International Conference on International Conference on Machine Learning, pages 833–840. Omnipress, 2011.

[18] Melissa Romeo and Max Diem. Correction of dispersive line shape artifact observed in diffuse reflection infrared spectroscopy and absorption/reflection (transflection) infrared micro-spectroscopy. Vibrational Spectroscopy, 38(1-2): 129–132, 2005.

[19] Nitish Srivastava, Geoffrey Hinton, Alex Krizhevsky, Ilya Sutskever, and Ruslan Salakhutdinov. Dropout: a simple way to prevent neural networks from overfitting. The Journal of Machine Learning Research, 15(1):1929–1958, 2014.

[20] Pascal Vincent, Hugo Larochelle, Yoshua Bengio, and Pierre-Antoine Manzagol. Extracting and composing robust features with denoising autoencoders. In Proceedings of the 25th international conference on Machine learning, pages 1096–1103. ACM, 2008.

[21] Pascal Vincent, Hugo Larochelle, Isabelle Lajoie, Yoshua Bengio, and Pierre-Antoine Manzagol. Stacked denoising autoencoders: Learning useful representations in a deep network with a local denoising criterion. Journal of machine learning research, 11(Dec):3371–3408, 2010.

[22] Yuanxin Xi and David M Rocke. Baseline correction for nmr spectroscopic metabolomics data analysis. BMC bioinformatics, 9(1):324, 2008.

[23] Hesham K Yosef, Sascha D Krauß, Tatjana Lechtonen, Hendrik Jutte, Andrea Tannapfel, Heiko U Kafferlein, Thomas Bruning, Florian Roghmann, Joachim Noldus, Axel Mosig, et al. Noninvasive diagnosis of high-grade urothelial carcinoma in urine by raman spectral imaging. Analytical chemistry, 89(12): 6893–6899, 2017.

[24] John R Zech, Marcus A Badgeley, Manway Liu, Anthony B Costa, Joseph J Titano, and Eric Karl Oermann. Variable generalization performance of a deep learning model to detect pneumonia in chest radiographs: A cross-sectional study. PLoS medicine, 15(11):e1002683, 2018.

[25] Zhi-Min Zhang, Shan Chen, and Yi-Zeng Liang. Baseline correction using adaptive iteratively reweighted penalized least squares. Analyst, 135(5):1138–1146, 2010.

