## Supplement A for "Deep representation learning for domain adaptatable classification of infrared spectral imaging data"

#### Supplementary Material

Arne P. Raulf, Joshua Butke, Claus Küpper,
Frederik Großerueschkamp, Klaus Gerwert, Axel Mosig

#### Contents

**S.1 Regularizing Autoencoders** **2**

**S.2 Validation of resulting segmentations** **2**

**S.3 Implementation Details** **3**

**Supplementary Figures and Tables** **3**

#### List of Supplementary Tables

1 Overview of supervised classifiers . . . . . . . . . . . . . . . . . . . . 6
2 Overview of data sets . . . . . . . . . . . . . . . . . . . . . . . . . . . 12

#### Supplement S.1 Regularizing Autoencoders

We follow the notion from the main paper, where an autoencoder is constituted as a neural network that represents a mapping $A\colon \mathbb{R}^d \to \mathbb{R}^d$ which, in its most basic form, involves one hidden layer with $M < d$ neurons.

An autoencoder can be seperated into an encoder $f\colon \mathbb{R}^d \to \mathbb{R}^M$ and a decoder $g\colon \mathbb{R}^M \to \mathbb{R}^d$, so that $A = g \circ f$. We will also refer to this type of autoencoder as a $(d, M)$ autoencoder. Assuming both $f$ and $g$ are two-layered feed-forward networks without hidden layers, we can write

$$\begin{aligned} f_\theta(x) &= Wx + b \quad \text{and} \\ g_{\theta'}(x) &= W'x + b' \quad , \end{aligned}$$

where $\theta := (W, b)$ and $\theta' := (W', b')$ are weight matrices and bias vectors constituting the parameters of the autoencoder. The autoencoder essentially generalizes principal component analysis and non-negative matrix factorization when trained under mean squared error. It has been shown [4] that adding the Frobenius norm of the Jacobi matrix of $g \circ f$ at $x$ as a penalty

$$L(x) := \|x - g(f(x))\| + \|J_F(g \circ f)x\|$$

achieves the desired regularizing effect. Our approach is based on the conjecture that the regularizing effect of stacked autoencoders can seperate the background artifacts from infrared microscopic pixel spectra, and thus learn a suitable preprocessing function. The complexity of the preprocessing function to be learnt in fact poses another essential problem for neural network approaches. Although in principle, a neural network may approximate a large class of almost arbitrary function [3], attempts to learn complex functions in practice tend to result in overfitting despite recent advances [5]. To adress this, different approaches towards *regularization* have been developed, which aim to reduce the complexity of the function to be learned, assuming that learning a less complex and thus more sparse description is more plausible than a very complex one.

#### Supplement S.2 Validation of resulting segmentations

To assess and compare the outcome of classifiers **pt-MLP**, **MLP** and **tl-MLP** against the respective ground truth, we employed accuracy as a validation measure. Whenever classifiers for FFPE tissue were validated on fresh tissue, any output class available in **RF2** (fresh tissue) but not in **RF** (FFPE tissue) (e.g. *mucinous cancer*, see Supplementary Fig. 3) was treated as background. Conversely, any pixel classified into a class provided by classifier **RF** but not available in ground truth provided by **RF2** (e.g. *slime*, see Supplementary Fig. 3) was counted as falsely classified. Similarly, classifiers **RF** and **RF2** require a (relatively small) number of pixels with low signal quality to be masked out. As no ground truth is available for these masked spectra, they were not taken into account for determining accuracies.

#### Supplement S.3 Implementation Details

We used stochastic gradient descent optimization for training the individual autoencoders during unsupervised pre-training in order to obtain the stacked autoencoder. We used a batch size of 500 and trained 2000 epochs at each level using a learning rate of $0.003$ and a sigmoidal activation function. Autoencoders were intitalized following [2].

For all supervised training tasks, we employed *RMSProp* optimization with a batch size of 500 and running up to 15000 epochs with a learning rate of $0.001$, ReLu activation function, and treating the the last 3 of the 6 layers as dropout layers with a dropout rate of $50\%$. Any layers that have not been pre-trained using the stacked autoencoder were initialized following [2].

All classifiers were implemented in Python using the *Theano* library for deep learning [1].

#### Supplementary Figures and Tables

*(see figures and tables on separate subsequent pages)*

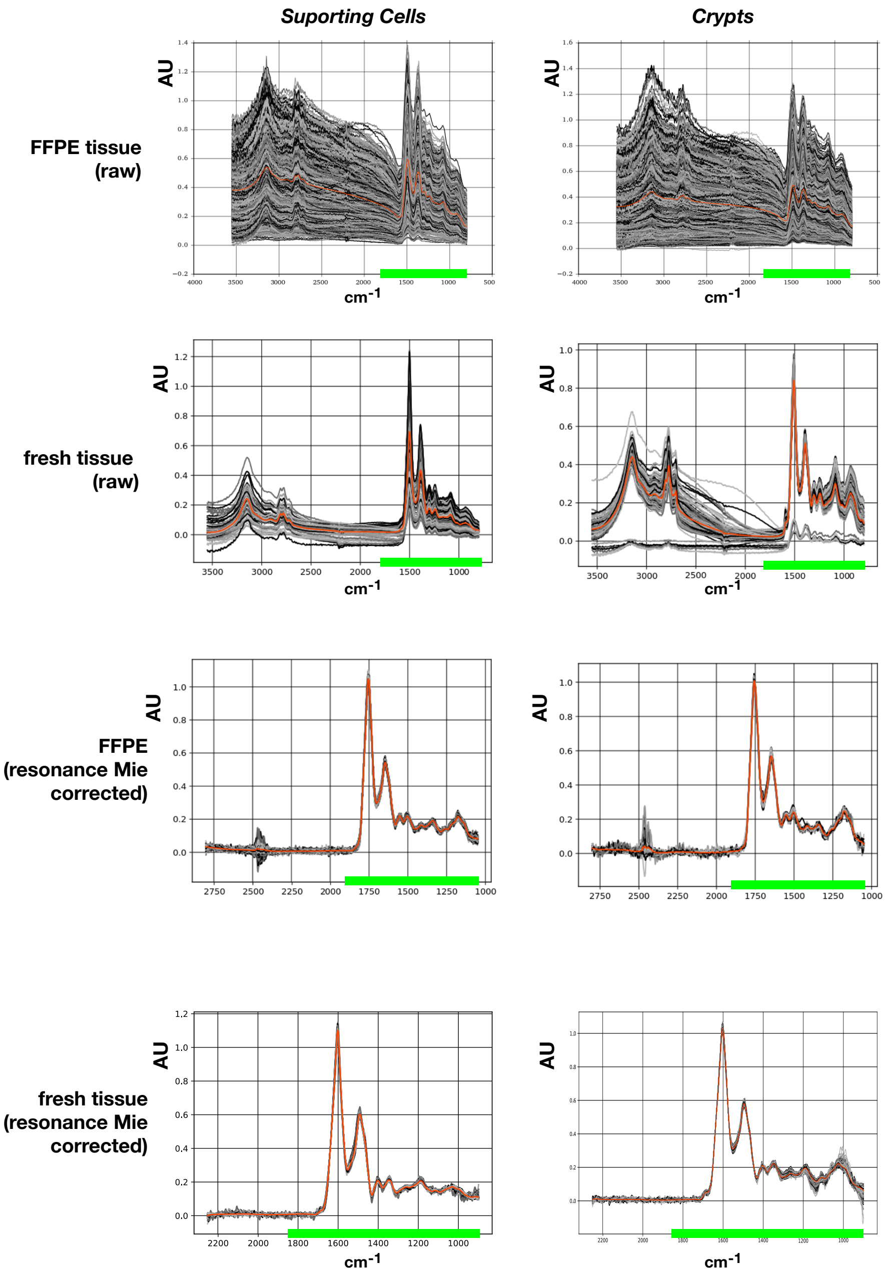

Supplementary Figure 1. Spectral Differences between FFPE-embedded tissue and fresh tissue before (*raw*) and after resonance Mie correction. Differences are shown for two spectra from two representative tissue components (supporting cells and crypts). Spectra have been drawn from classification results of classifiers **RF** and the transfer-learned **pt-MLP** for fresh tissue. Individual spectra have been plotted in randomly assigned gray scales; the average spectrum is plotted in orange. The green bar indicates the region between 950 $cm^{-1}$ and 1815 $cm^{-1}$ which has been utilized during all supervised training steps (classifiers **pt-MLP**, **MLP** and transfer learning from FFPE to fresh tissue). Uncorrected raw spectra naturally exhibit a high variance that is particularly strong for FFPE spectra. Resonance Mie correction obviously has an effect of eliminating variance in the spectra.

### References

[1] James Bergstra, Olivier Breuleux, Frédéric Bastien, Pascal Lamblin, Razvan Pascanu, Guillaume Desjardins, Joseph Turian, David Warde-Farley, and Yoshua Bengio. Theano: A cpu and gpu math compiler in python. In *Proc. 9th Python in Science Conf*, volume 1, 2010.

[2] Xavier Glorot and Yoshua Bengio. Understanding the difficulty of training deep feedforward neural networks. In *Proceedings of the thirteenth international conference on artificial intelligence and statistics*, pp. 249–256, 2010.

[3] Kurt Hornik. Approximation capabilities of multilayer feedforward networks. *Neural networks*, 4(2):251–257, 1991.

[4] Salah Rifai, Pascal Vincent, Xavier Muller, Xavier Glorot, and Yoshua Bengio. Contractive auto-encoders: Explicit invariance during feature extraction. In *Proceedings of the 28th International Conference on International Conference on Machine Learning*, pp. 833–840. Omnipress, 2011.

[5] Nitish Srivastava, Geoffrey Hinton, Alex Krizhevsky, Ilya Sutskever, and Ruslan Salakhutdinov. Dropout: a simple way to prevent neural networks from overfitting. *The Journal of Machine Learning Research*, 15(1):1929–1958, 2014.

| **Classifier** | *Description* | *Purpose* |
|---|---|---|
| Classifier pt-MLP | Pretrained multilayer perceptron; pretrained using the unsupervised stacked contractive autoencoder, followed by supervised fine-tuning. | *Main contribution.* Pretraining claims to learn a representation of infrared spectra which is favourable towards classification and generalization of these classifiers. |
| Classifier MLP | Conventional multilayer-perceptron; uses dropout layers, but no pre-training for regularization. | Reference classifier to compare with the pre-trained classifier **pt-MLP**. |
| Classifier tl-MLP | Obtained by transfer learning, starting from classifier **pt-MLP** and using spectra with corresponding ground truth labeling from fresh tissue as training data. | *Main contribution.* Assess the generalization strength of classifier **pt-MLP** and of the underlying unsupervised pretraining. |
| Classifier RF | Previously established random forest classifier to identify tissue components, including tumor regions, of FFPE-embedded colon tissue. | Provide ground truth to train classifier **pt-MLP**. |
| Classifier RF2 | Previously established random forest classifier to identify tissue components, including tumor regions, of fresh colon tissue. | Provide ground truth for transfer learning to train classifier **tl-MLP**. |

Supplementary Table 1. Overview of supervised classifiers used throughout the paper

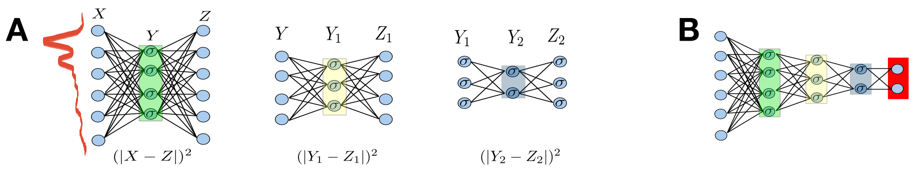

Supplementary Figure 2. Schematic illustration of a stacked autoencoder. Panel **A** displays the cascade of autoencoders, where each layer is trained to encode the encoded representation of the previous layer. The resulting autoencoders can be cascaded as illustrated in panel **B** on the right. The last layer represents the output layer, which is added during supervised fine-tuning on labelled training data. The illustration is schematic only, and the autoencoder established in our contribution involves an input layer with input neurons for 450 wavenumbers of an infrared spectrum, an output layer involving 100 output neurons, and six hidden layers of sizes $M_1, \ldots, M_6 = 450, 900, 450, 100, 100, 100$.

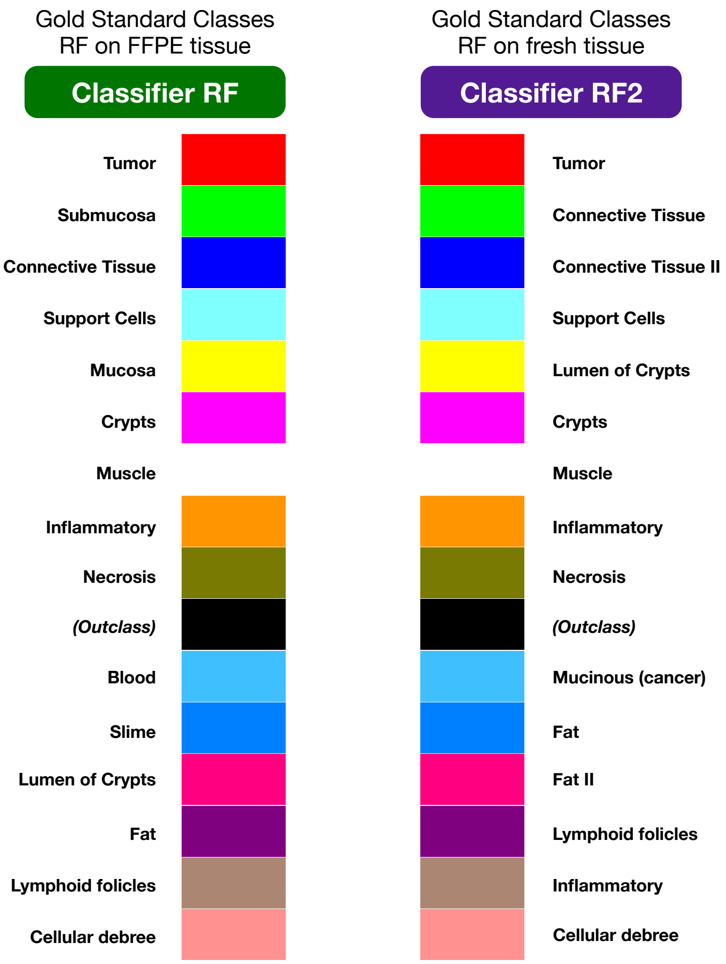

Supplementary Figure 3. Classes of the two classifers for the gold standard obtained from random forest classifiers for FFPE tissue *left* and fresh tissue *(right)*.

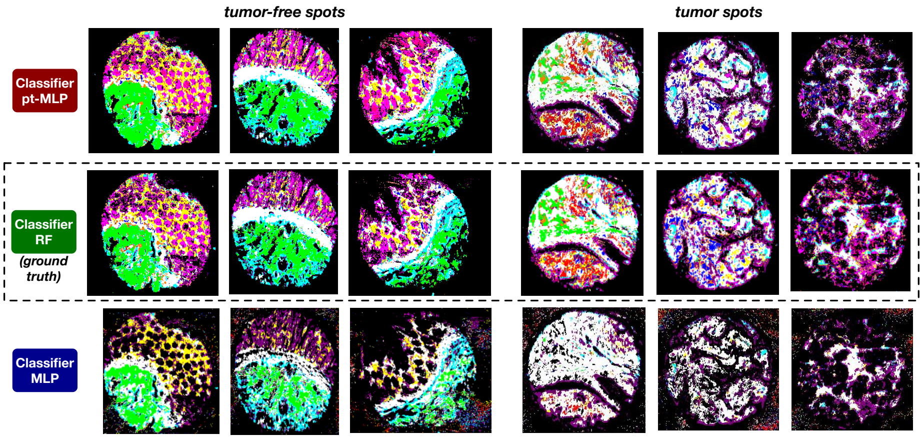

Supplementary Figure 4. Segmentation results of six validation spots comparing the results of classifiers **pt-MLP** and **MLP** with the ground truth obtained from classifier **RF**.

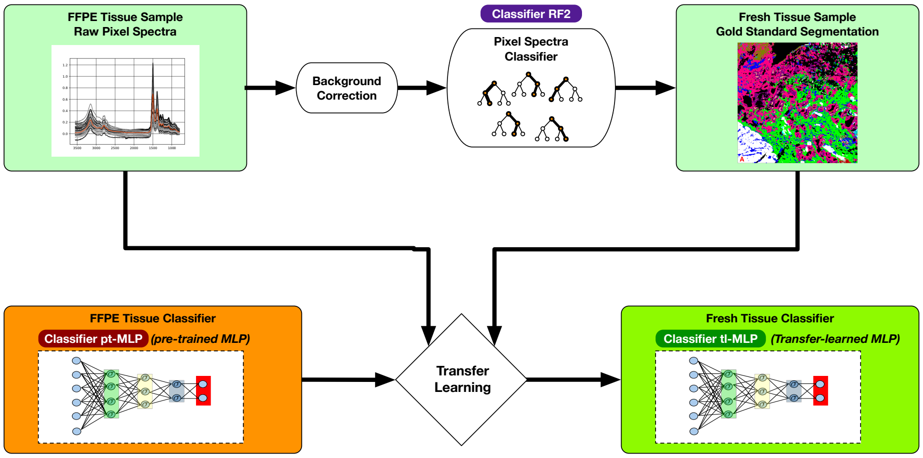

Supplementary Figure 5. Schematic overview of transfer learning. We use classifier **pt-MLP** as a basis along with uncorrected raw spectra from fresh tissue samples, and ground truth tissue class labels assigned to these spectra using a previously established classifier for background corrected spectra from fresh colon tissue.

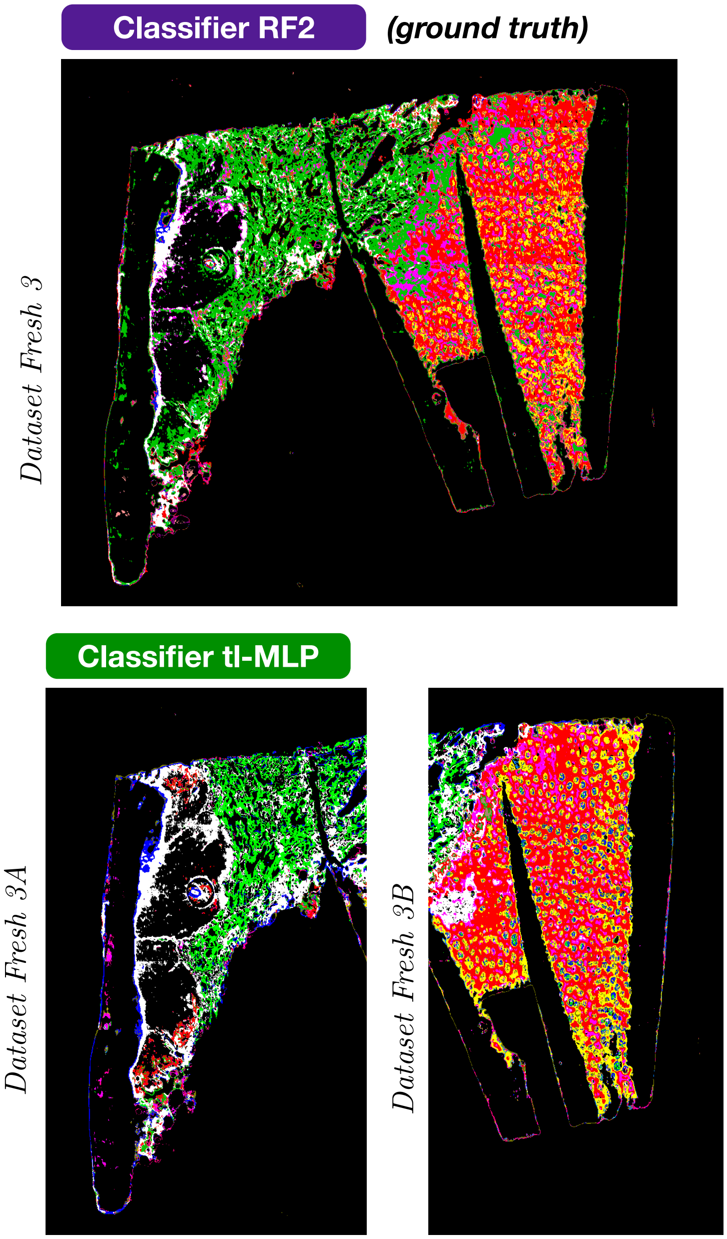

Supplementary Figure 6. Transfer Learning results on data set *Fresh 3*. Classifier **tl-MLP** achieved accuracies of 72% on data set *Fresh 3A* (left) and 80% on data set *Fresh 3B* (right).

| **ID** | **Preparation** | **Type** | **Usage** |
|---|---|---|---|
| *CO722* | FFPE | tissue microarray (Biomax) | unsupervised pre-training |
| *CO1002b* | FFPE | tissue microarray (Biomax) | training and validation of classifier **pt-MLP** |
| *Slide 1* | fresh-frozen | whole-slide | ground truth for transfer-learning classifier **tl-MLP** |
| *Slide 2* | fresh-frozen | whole-slide | validation of classifier **tl-MLP** |
| *Slide 3* | fresh-frozen | whole-slide | validation of classifier **tl-MLP** |

Supplementary Table 2. Overview of data sets used in this study.
